# The evolution of investment in innate-like and diversified T cell receptors across development

**DOI:** 10.1101/2025.09.04.674346

**Authors:** Reese A. Martin, Anna E. Savage, Ann T. Tate

## Abstract

New insights into the diversity of lymphocyte functions challenges previous dogma about the rigid divide between innate and adaptive immunity. While T cells with canonically diversified receptors are crucial for recognizing novel antigens, other T cell lineages express innate-like receptors that recognize conserved molecular patterns. The relative frequency of innate-like to diversifying T cell receptors (iTCRs: dTCRs) varies greatly across vertebrate species and across ontogeny within species. These within-species dynamics can potentially be explained by developmental constraints on immunity, pathogen diversity and exposure, or by trade-offs associated with specificity. To better understand how these factors shape T cell repertoires, we constructed an agent-based model of TCR evolution inspired by the diversity of ontogenic life histories in amphibians but applicable to an array of vertebrate species. Our model features two life stages with distinct parasite populations and life history costs. The model predicts that changes in ontogeny (stage duration, T cell maturation time) and environmental factors (parasite diversity, parasite complexity) exert drastic effects on the stage structure of T cell investment strategies. A better understanding of the evolutionary pressures that shape TCR diversity will provide new insights into lymphocyte evolution and immune investment across organismal development.

## Introduction

T lymphocytes express receptors (TCRs) that are collectively capable of recognizing a high diversity of novel peptides associated with infection and abnormal self. T cells destined to bear these diversified receptors go through somatic recombination to generate a novel receptor. They then endure positive and negative selection in the thymus to maximize the probability that the receptor complex will achieve robust activation if it does recognize a foreign peptide, while at the same time avoiding activation due to healthy self-peptides. These diversified TCRs (dTCRs) compose greater than 90% of the TCR repertoire in adult humans (Girardi 2006) and – alongside B cells – are traditionally hailed as a cornerstone of the vertebrate adaptive immune system. Not all T cells bear these dTCRs, instead expressing a more predictable class of innate-like TCRs (iTCRs) (Van Kaer et al. 2022). These cells include invariant (NK)T cells, mucosal-associated invariant T cells (MAITs), and some γδT cells, all of which are united in expressing a TCR that functions akin to pattern-recognition receptors of the innate immune system rather than a dTCR that must undergo the gauntlet of selection and MHC restriction (Li and Wu 2021).

The existence of these distinct classes of TCRs raises a question: why bother creating T cells bearing these innate-like receptors rather than just relying on the innate immune system to recognize conserved motifs? After all, an activated T cell lineage can stimulate a substantial inflammatory response and promote damage to the host through autoimmunity and immunopathology if the response is not finely controlled. One major clue comes from the relative ratios of innate-like versus diversified TCRs across organismal development from embryonic hematopoiesis to maturity. In vertebrates as evolutionarily distant as frogs, mice, and humans, innate-like TCRs and associated T cell subsets dominate in early life and give way to a primarily diversified repertoire later in life, although these ontogenetic patterns vary from taxon to taxon (Siegrist and Aspinall 2009; Edholm et al. 2013; Papadopoulou et al. 2020). Notably, even the smallest vertebrates successfully mount competent immune responses using dTCRs, suggesting that the transition from innate-like to diversifying T cells is not explained by body size or the physical (in)ability to support hundreds of billions of T cells (Giorgetti et al. 2021).

In humans, γδT cells, which do not rely on MHC and recognize conserved phosphoantigens and lipids, expand during fetal development and reach their maxima in peripheral T cell populations soon after birth, giving way to the dominance of diversified αβT cells by early childhood (Clark and Thomas 2020). γδT cells with innate-like functions do not just disappear, however; they remain in high numbers within mucosal tissues (Kang et al. 2023). The tadpole stage of the frog *Xenopus laevis* is similarly dominated by innate-like αβTCRs with low variability in both gene segment use and nucleotide diversity, whereas the adult stage exhibits a full repertoire of dTCRs (Robert and Edholm 2014). In addition to receptor diversity, these tadpole and adult T cell subsets differ in their proliferative and inflammatory potential. Tadpoles exhibit a less inflammatory and more tolerogenic T cell response to common pathogens like ranaviruses and *Mycobacterium marinum* (Rhoo et al. 2019). While this is associated in some instances with increased susceptibility to infection-induced mortality, it may also dampen the accumulation of pathological damage that would impair the reproductive fitness of subsequent life stages (Metcalf et al. 2017; Oyesola et al. 2024).

Life history trade-offs thus provide one explanation for the dynamic nature of TCR diversity across ontogeny, but developmental and ecological factors may also contribute to these changes. iTCRs are likely honed over evolutionary time to recognize motifs associated with predictable enemies, as with MAIT cells targeting conserved bacterial and fungal ligands (Nel et al. 2021), whereas diversifying TCRs provide a means to respond to novel pathogen epitopes. Thus, stage-specific parasite diversity and exposure rates in the wild could drive selection for different TCR strategies in different life stages. The differences may also come down to developmental constraints. For example, the tadpole thymus expresses terminal deoxynucleotidyl transferase (TdT) at lower levels than the adult, limiting the addition of random nucleotides to variable regions (Paiola et al. 2023). Diversified T cell maturation in the thymus also involves a substantial investment of time and energy that fast-maturing organisms may not benefit from. For example, if it takes two weeks for a thymocyte to undergo positive and negative selection, migrate to a lymph node, find its cognate antigen, and achieve activation it would not make sense for an organism that completes its larval stage in ten days to investment in diversified T cell mediated immunity.

This all assumes, of course, that organisms are capable of deploying different strategies in different life stages. If the genetic or physiological basis of an immune trait is pleiotropic with other stage-specific traits, for example, a change in immune expression to optimize defense in one life stage could force covarying changes to other traits to the detriment of organismal fitness. Antagonistic pleiotropy is one example of the myriad processes that can exact a decoupling cost, the penalty to fitness that arises when organisms express two or more distinct phenotypic regimes across different life stages. Decoupling costs share many conceptual similarities with the costs of phenotypic plasticity (Relyea 2002), where decoupled traits are akin to highly plastic ones.

Metamorphosis may alleviate some types of decoupling costs, as it allows remodeling of the body plan between life stages, providing a means to decouple traits and optimize the response to selection pressures facing each stage individually (Robert et al. 1997). However, metamorphic species may face high metabolic requirements to remodel organs and genomic constraints on regulation and expression necessary to organize multiple complex phenotypes. The trade-offs associated with these types of decoupling costs may even explain the coexistence of metamorphic taxa and direct developers like *Callulina sp.* (warty frogs) along the anuran (frog) lineage, facultative neoteny in multiple salamander clades (Denoël et al. 2005), and the rise of neoteny in insect lineages otherwise capable of complete metamorphosis (Mcmahon and Hayward 2016). Here metamorphosis offers species the ability to decouple trait performance across life stages (Johansson et al. 2010) with a trade-off centered on the energetic capital to fuel the metamorphic transition, and metamorphic stages (e.g. insect pupae) being particularly susceptible to predation.

Despite the multitude of anecdotes describing distinct age-specific patterns of innate-like and diversified T cell prevalence in different vertebrate taxa, we lack a general theory for predicting when organisms should invest in one versus the other. A second open question relates to the degree to which an organism should invest in their diversifying immune response. First and foremost, are predictions for the evolution of innate vs adaptive immunity generally transferrable to the problem of T cell receptors, or does the unique inflammatory and dynamical profiles of iTCRs put them in a distinct conceptual class? Should early life stages always skew investment toward iTCRs, or could parasite ecology, developmental factors, or decoupling costs promote a diversity of strategies within and among species? Drawing on the diversity of amphibian life histories, where larval stages, for example, can span 5-75% of the time until sexual maturity (Table 1), we developed an agent-based evolutionary model to study how environmental and ontogenic factors shape the balance between iTCRs and dTCRs in host immunity.

**Table 1.** The diversity of stage length and assayed microbiota across anuran species. Note: OW = Overwinters, FPD = Faith’s Phylogenetic Diversity, LogGB = Log (Gram-negative bacterial counts)/g/ml, ASV = amplicon sequence variants, m = male, f = female

| Species | Larval lifespan (months) | Adult lifespan (years) | Age of sexual maturity (years) | Larval microbial diversity | Adult microbial diversity | Microbial taxa | Metric | references |
| --- | --- | --- | --- | --- | --- | --- | --- | --- |
| *Anaxyrus boreas*<br>(Western Toad) | 1 - 1.5 | 9 -12 | 4 - 6 | 2 | 5 | skin bacteria | Shannon alpha diversity | (Prest et al. 2018) |
| *Anaxyrus americanus*<br>(American Toad) | 1 - 2 | Up to 10 | 2 - 3 | 22.5 | 21 | skin bacteria | FPD | (Jones et al. 2025) |
| *Bufo terrestris*<br>(Southern Toad) | 1 - 2 | 10 | 2 - 3 | 5.4 | 5.9 | gut bacteria | LogGB | (Fedewa 2006) |
| *Hyla versicolor*<br>(Gray Treefrog) | 1.5 - 2 | 7 - 9 | 2 | 11 | 16 | skin bacteria | FPD | (Jones et al. 2025) |
| *Pseudacris crucifer*<br>(Spring Peeper) | 1.5 - 3 | 3 - 4 | 1 | 3.7 | 5.45 | gut bacteria | LogGB | (Fedewa 2006) |
| *Bombina orientalis*<br>(Oriental Fire-bellied Toad) | 2 | Up to 20 | 2 - 4 | 145 | 105 | skin bacteria | Chao 1 index | (Bataille et al. 2018) |
| *Rana cascade*<br>(Cascades Frog) | 2 | 5 - 7 | 2 - 4 | 18 | 33 | skin bacteria | FPD | (Kueneman et al. 2014) |
| *Rana sylvatica*<br>(Wood Frog) | 2 - 4 | 3 - 5 | 1 - 2 (m),<br>2 - 3 (f) | 17 | 14.5 | skin bacteria | FPD | (Jones et al. 2025) |
| *Notophthalmus viridescens*<br>(Eastern Newt) | 2 - 5 | 9 - 15 | 1 - 3 | 110 | 105 | gut bacteria | ASV | (Fontaine et al. 2021) |
| *Dryophytes (Hyla) japonicus*<br>(Japanese Tree Frog) | 2 - 10<br>(OW) | 6 - 11 | 3 - 4 | 20 | 55 | gut bacteria | FPD | (Park et al. 2023) |
| *Triturus marmoratus*<br>(Marbled Newt) | 2 - 10 | 15 | 5 | 252.7 | 359.7 | skin bacteria | Chao 1 index | (Sabino-Pinto et al. 2017) |
| *Ambystoma mavortium*<br>(Western Tiger Salamander) | 2 - 12 | 2 - 3 | 4 - 5 | 9.5 | 5.5 | Skin fungi | Hill's diversity index | (Goodwin et al. 2022) |
| *Lissotriton boscai*<br>(Bosca's Newt) | 2 - 12 | 8 | 2 - 4 | 349.9 | 509.2 | skin bacteria | Chao 1 index | (Sabino-Pinto et al. 2017) |
| *Rana pipiens*<br>(Northern Leopard Frog) | 3 - 6 | 5 - 8 | 1 - 3 | 26 | 19 | gut bacteria | FPD | (Kohl et al. 2013) |
| *Notophthalmus peristriatus*<br>(Striped Newt) | 6 - 9 | 12 - 15 | 1 | 8 | 32 | skin bacteria | FPD | (Hartmann et al. 2023) |
| *Quasipaa spinosa*<br>(Giant Spiny Frog) | OW | 2 - 4 | 1 | 344 | 318 | skin bacteria | ASV | (Hou et al. 2024) |

Organisms in this model go through distinct larval and adult life stages and are exposed to different parasite populations in each. Host immunity is represented by a TCR repertoire comprising iTCRs and dTCRs. Hosts are born with a selection of iTCRs well-matched to a subset of the larval parasite population, and life-stage specific dTCR investment drives the creation dTCRs. Organisms reproduce in the adult life stage, and their relative fitness is governed by a function that balances the metabolic and pathological costs of creating and deploying each TCR subset against the pathological costs of parasite burden (fig. 1). We used this model to determine how relative changes in stage length, immune costs, and decoupling costs between adult and larval dTCR investment could act together to shape the balance of iTCRs and dTCRs in host TCR repertoires. Taken together, our results suggest that decoupling costs are an important piece of the puzzle for understanding ontogenic investment patterns and for explaining natural variation in investment within and across species.

**Figure 1:**
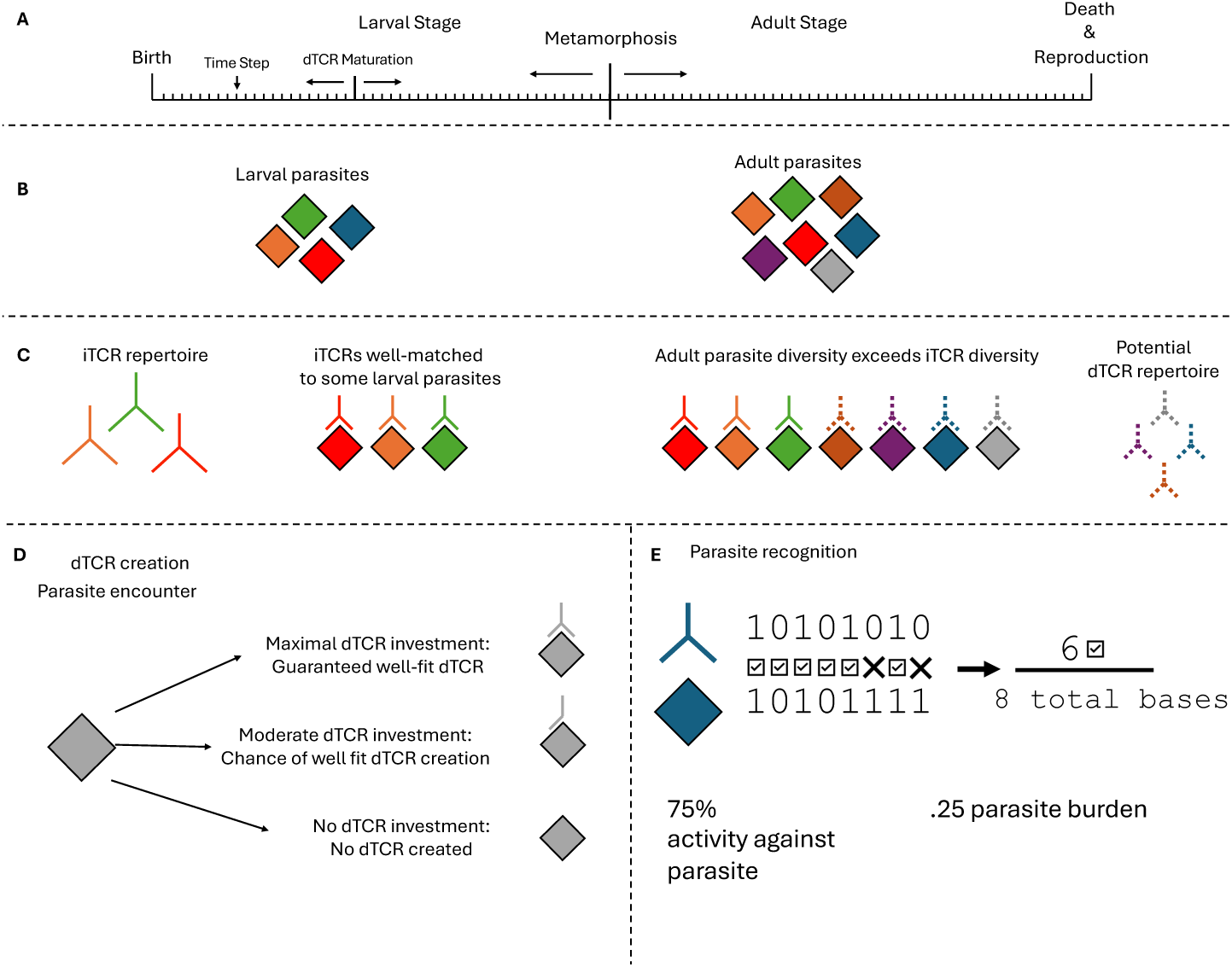
Visual depiction of key model components. **A)** Hosts undergo a 100-timestep lifespan where they face a chance of parasite exposure at each timestep. Hosts begin life as larvae before transitioning to an adult stage. After a specified number of timesteps, the thymus and first thymocytes mature and they gain access to diversified immunity (labeled dTCR maturation). During metamorphosis, hosts switch from their larval to adult dTCR investment strategy, and their potential parasite milieu switches to the adult set. **B)** Each color represents a different species of parasite to which hosts may be exposed; here adult parasite diversity exceeds larval parasite diversity and larval parasites are a subset of the adult population, but that assumption can be relaxed. **C)** Larvae are born with an iTCR repertoire that is constant across life stages. Each iTCR can be matched against a parasite, but in cases where parasite diversity exceeds iTCR diversity, hosts can generate dTCRs (dashed lines). iTCRs are inherited across generations but dTCRs are not. **D)** The level of dTCR investment determines the relative probability of a partial or full match between dTCR and novel parasite ligand. If a newly created dTCR exceeds the best fit of all current TCRs against the parasite, then it is added to the host TCR repertoire. **E)** TCR-Parasite Fit. Each parasite and TCR comprise a series of 1s and 0s (bases). If a TCR base matches a parasite base at the same position, that contributes to the TCR’s fit against the parasite. The parasite burden (fitness cost due to parasite infection) is 1-proportion TCR-Parasite Fit.

## Methods

### Model overview and predictions

We developed an agent-based modeling framework to study host immune investment strategies. We simulated host life from birth to the age of first reproduction, with all hosts starting each generation in the larval stage and ending in the adult stage. To ensure that dTCR investment in the final generation of the simulation was not due to initial condition biases, the first generation of each simulation begins with a population of hosts that have no investment in dTCR-mediated immunity (see Figs. S1,S2 for simulations where hosts started with intermediate or maximal investment). For each host in each generation, larval and adult life stages have independent dTCR investment strategies and semi-independent parasite populations. At the end of each generation, hosts reproduce and die in a fitness-weighted manner (see methods section *Evolutionary Simulations 3. Fitness* for details concerning host fitness calculations).

Optimizing host reproductive fitness requires balancing immune investment against the costs of parasitism (as described by *Eq. 2* below). Biologically, both immune activity and parasite burden can impose metabolic and pathological costs to host fitness, while dTCR investment incurs additional metabolic costs relative to iTCRs associated with culling 95% of thymocytes during selection and the risk of autoimmunity (Merkenschlager et al. 1997; Song et al. 2024). Fitness costs due to immunity were composed of the cumulative lifetime immune action taken by hosts, dTCR investment from both life-stages, and the decoupling costs for mounting differential dTCR responses between life stages. Lifetime parasite burden, which captured the mismatch between the host’s TCR repertoire and the parasite, represented the parasite’s contribution to reduction in host fitness.

New mutations arose in host dTCR immune investment and iTCR repertoire during reproduction (See *Evolutionary Simulations*: 5. Reproduction). Parasite populations are kept constant in each simulation, reflecting one-sided evolutionary dynamics consistent with large communities that share generalist parasites (Rabajante et al. 2015) rather than coevolution. Table S1 contains model parameters, their default values, and the biological/environmental trait they reflect.

We explored several questions related to the evolution of dTCR investment that required toggling different parameters individually and in combination. We first wanted to ground-truth our model against well-established theory on the general evolution of immune investment (Mayer et al. 2016; Metcalf et al. 2017; Oyesola et al. 2024). This means that our results should adhere to the following basic predictions:

1. dTCR investment should increase as parasite diversity increases, especially when parasite diversity exceeds the capacity of innate-like receptor recognition. If hosts are never exposed to a higher parasite diversity than their iTCR repertoire capacity, then they should not evolve dTCRs because dTCRs are more costly than iTCRs.
2. dTCRs should not evolve if their cost in maturation, deployment, or both, far exceeds any benefit in resistance.

Stage structure could add complexity to these predictions, however. Most notably, larvae and adults could differ in the diversity of the parasites to which they are exposed and the types of costs they incur from immune investment. For example, larva may face trade-offs between development and immunity (Wittkopp and Beldade 2009; Alzaid et al. 2016) while trade-offs between immunity and reproduction impose costs on adults (McKean and Nunney 2005; Schwenke et al. 2016). In addition, there may be a benefit to “anticipating” adult parasite diversity by beginning dTCR maturation in the larval stage, since dTCRs are not purged during metamorphosis (Robert et al. 1997). Finally, there could be a structural or genetic “decoupling cost” to deploying different strategies in different life stages, which could lead to suboptimal investment in one or both life stages. Thus, we made the following predictions:

1. If the larval stage is too short relative to the adult stage or to the time of dTCR maturation, this should preclude the evolution of larval dTCR investment.
2. If there is a large decoupling cost to immunity, immune investment in both stages will mirror optimal investment in the stage with the larger parasite diversity regardless of the optimal investment strategy in the other stage.

### TCR-Parasite Fit and dTCR production

Hosts and parasites interact using the host TCR repertoire and parasite base sequence (fig. 1E). In this model, parasites and TCRs comprised sequences of 1s and 0s (referred to as bases) of a static length that was specified at the start of each simulation (default: 20). The TCR-Parasite Fit was the percentage of TCR bases that matched the corresponding parasite bases (fig. 1E). TCR-Parasite Fit determined the burden of the infection on hosts. A perfect fit, where all bases matched, resulted in no burden, while a perfect mismatch resulted in a maximal parasite burden of 1.0.

The host TCR repertoire includes both genetically encoded iTCRs and any dTCRs produced during the course of its life. Hosts produced dTCRs in response to infection when the following two conditions were met: 1) the host evolved investment in dTCR immunity for the focal life stage, and 2) the infection occurred after the dTCR immune system matured. If the resulting dTCR had a higher percent match against the infecting parasite than all other TCRs, the dTCR was added to the host’s TCR repertoire and was available for their remaining lifespan. Otherwise, the dTCR was discarded and the best fit TCR repertoire was used to fend off the infection. In both cases the hosts paid the dTCR fitness cost. In simulations where a maturation delay was added to dTCR creation, the generated dTCR was not added to the host TCR repertoire until the requisite number of timesteps had passed. The percent match of generated dTCRs was determined using *Eq. 1*:

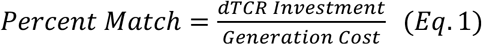

where *dTCR Investment* is the host’s dTCR investment and *Generation Cost* is a parameter controlling the amount of investment needed to produce a perfectly matched dTCR.

#### Parasite Diversity

Larval and adult parasite population diversity was specified at the start of each simulation. Each parasite was generated as a random 20-base sequence of 1s and 0s. Larval parasites were generated first, then an additional *n* parasites were added to that subset to create the adult parasite population. In scenarios where larval parasites outnumbered adult parasites, a subset of the larval parasite population was selected to generate the adult parasite population. In this way the smaller parasite population was always a subset of the larger parasite population.

### Evolutionary simulations

#### Initialization

at the start of each simulation, 500 hosts were generated along with a specified number of larval and adult parasites. Host iTCRs were then created with each iTCR matching perfectly against one larval parasite. If there were fewer larval parasites than iTCRs, the excess iTCRs were generated at random. Supplemental Table 1 contains the default values for the parameters used in these simulations.

Every generation cycled through steps 1-5 below; the next generation used the population derived in step 5 of the previous generation. Unless otherwise specified, a simulation consisted of a population of 500 hosts evolving through 1000 generations, and the results were the average of 100 simulations.

1. **Larval Stage:** The larval stage starts at timestep 1 and lasts a default 25 timesteps; however, variation in the length of the larval stage relative to total time to reproductive maturity is a key variable under investigation (e.g., Table 1). Infections were drawn from the pool of available larval parasites and occurred with a fixed probability per timestep (default: 0.05/timestep). If a host was infected, their current TCR repertoire was tested to find the best matching TCR for the infecting parasite. A host with no dTCR investment relied solely on iTCR defenses, otherwise dTCR creation, deployment, and determination of parasite burden was calculated as described above. We assumed that larvae with a poorly controlled parasite burden would die from disease before or during metamorphosis. Therefore, larvae with a parasite burden > 1 at the cusp of the adult transition were removed from the population, while the rest entered the adult stage.
2. **Adult Stage:** The adult stage lasted from the end of the larval stage to the end of host reproductive lifespan (100 total timesteps unless otherwise specified) at which point hosts died and reproduced in a fitness-weighted manner. In this stage hosts were exposed to parasites from the adult parasite population and dTCRs were generated according to the adult dTCR investment strategy. Infection burden was determined as in step 1.
3. **Fitness:** Host fitness was calculated using three components: 1) the cost of investment in dTCR maturation, 2) the costs associated with deployment of the immune response, and 3) costs accumulated from parasite burden. Fitness was evaluated using values accumulated over the entire lifespan. The effects of these individual components on organismal fitness were determined using Equation 2:

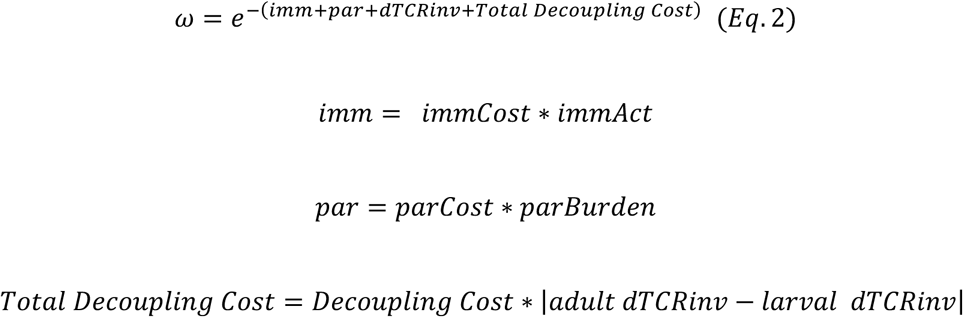

where *immCost* was a parameter modifying the cost of general immunity, *immAct* was the host’s cumulative immune activity, *parCost* was a parameter modifying the general cost of parasitism, *parBurden* was the host’s cumulative parasite burden, *dTCRinv* was the sum of dTCR investment from both host life stages, and *Decoupling Cost* was a parameter that determines the fitness cost for differential dTCR investment between life stages.
4. **Death:** Hosts died in a fitness-weighted manner, where a host’s chance of death was inversely proportional to its relative fitness in the population. Each host in the population was evaluated in this way, and there was no limit to the number of hosts that could die in a single generation.
5. **Reproduction:** following the removal of dead hosts from the population, hosts reproduced proportionally to their relative fitness. Hosts were selected at random for the chance to reproduce, and reproduction continued until the population returned to the initial population size. Hosts were not restricted to a single reproductive event per generation.

a. **Mutation:** Offspring were born into the next generation as direct copies of individual parents (no recombination) with the exception that dTCRs were not passed to offspring. During reproduction there was a small chance (p = 5e-3) that a mutation would occur. Mutations and their relative probabilities were as follows: 1) Delete an iTCR (p=0.25); 2) Generate a random iTCR (p = 0.25); 3) Increase or Decrease larval dTCR investment, (p = 0.25); 4) Increase or Decrease adult dTCR investment (p=0.25).

Following stage 5, the generation concluded, and step 1 began again with the new population.

### Phylogenetic analysis comparing larval stage length and microbial diversity

We were curious whether there was enough data in Table 1 to determine whether stage duration was indeed correlated to increased parasite diversity. We analyzed the relationship between larval stage length and microbial diversity (Supplemental Methods) and found a weak positive relationship between microbial diversity ratio (Larval: Adult) and larval stage length (fig. S3). However, we cannot be certain that overall microbial diversity is a reliable proxy for pathogen diversity and more investigation is needed before drawing detailed conclusions from this analysis. As such, we omitted more rigorous parameterization of the model based on these data, opting instead to present more general results with independent life-stage length and parasite diversity parameters.

## Results

### dTCR investment increases with parasite diversity and is sensitive to both stage length and decoupling costs

To confirm that our model recapitulated well-established predictions from evolutionary theory, we first simulated conditions where the percent of life as a larva was varied against the percent of life before the diversifying immune response matured, in the absence of any decoupling cost (fig. 2A,B; the asterisk in panel C shows the dTCR cost and adult parasite diversity used for these simulations). When varying the percent of life spent as a larva against dTCR maturation time, we observed that larvae only invested in dTCRs when larval lifespan was longer than the time for dTCR maturation (fig. 2A), while adults always invested in dTCR immunity (fig. 2B). Adding additional types of delays, like a delay between infection and dTCR creation, served to shift larval dTCR investment to the left and was effectively identical to increasing the dTCR maturation delay by the second delay term (fig. S2). Changing the initial dTCR investment conditions so that hosts started with intermediate or maximal investment provided similar results except that high initial investment reduced larval responsiveness to adult parasite diversity under different decoupling cost conditions (figs. S1, 2).

**Figure 2.**
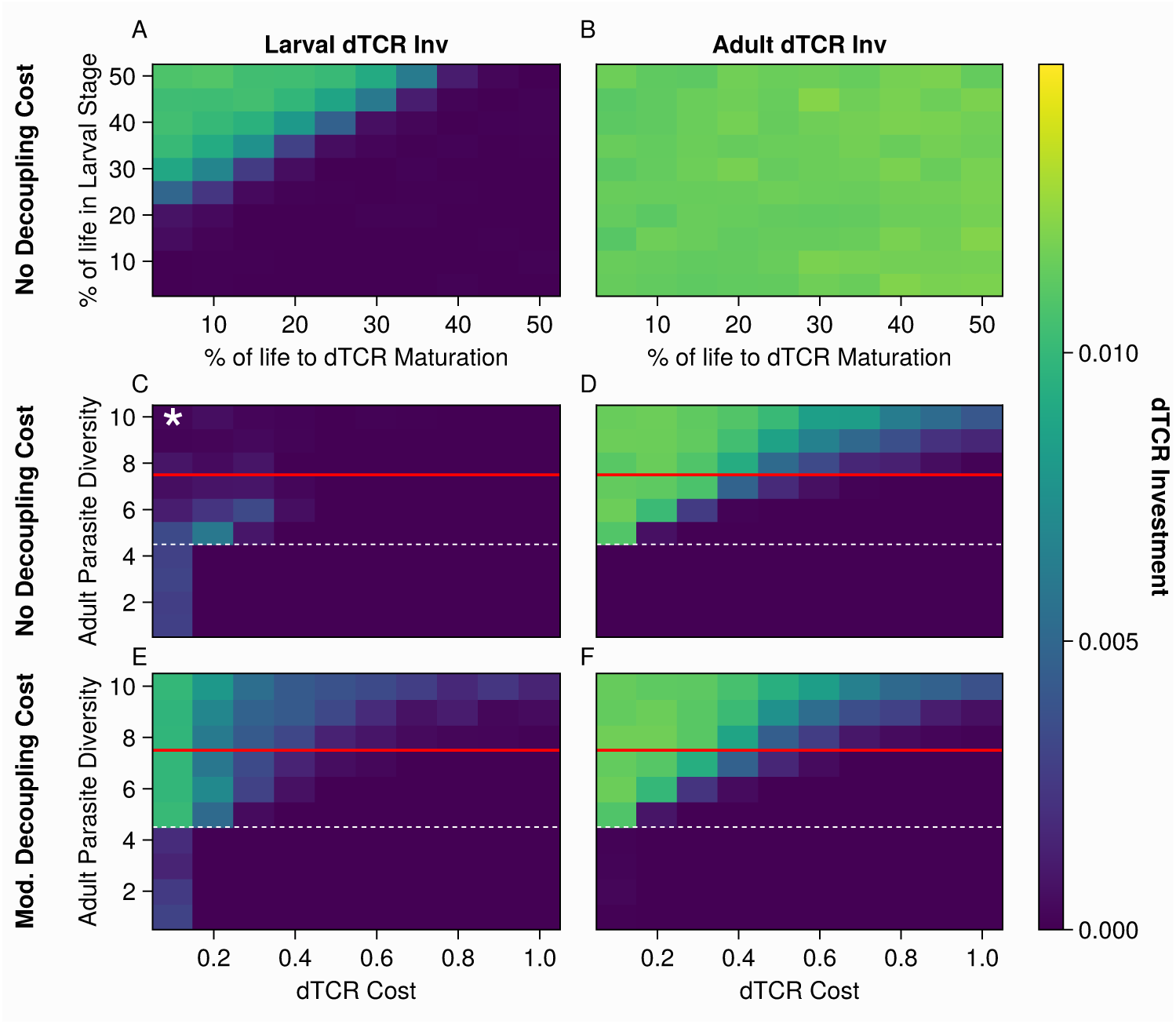
dTCR investment generally increases with higher parasite diversity and lower costs, but carry-over effects and the relative length of larval stage both influence optimal larval investment. Warmer colors indicate greater dTCR investment. A, B: y-axis shows the percent of life in the larval stage, x-axis shows the percent of life before the diversifying immune response matures and can be used to fend off parasites. C, D: No decoupling cost. y-axis shows adult parasite diversity; x-axis shows the cost associated with generating dTCRs. The white dashed line shows iTCR repertoire diversity and the solid red line shows larval parasite diversity. E, F: Moderate decoupling cost. The y-axis shows adult parasite diversity; x-axis shows the cost associated with generating dTCRs. The white dashed line shows iTCR repertoire diversity and the solid red line shows larval parasite diversity.

To test the prediction that parasite diversity and dTCR costs influence dTCR investment, the cost of diversifying immunity was varied against adult parasite diversity (fig. 2C-F). As adult parasite diversity increased, adult investment in dTCRs also increased, while increasing dTCR costs led to decreased dTCR investment (fig. 2C). Adults only began investing in their dTCR response after the adult parasite population exceeded the iTCR repertoire diversity (dashed line, fig. 2C-F). Larval dTCR investment peaked in the window where adult parasite diversity surpassed iTCR repertoire diversity but remained smaller than larval parasite diversity, which was constant throughout simulations (red line, fig. 2C-F). Hosts in this window of relative parasite diversities appear to invest in dTCR immunity as either adults or larvae, with very few investing in both stages.

Having established that the behavior of the model captures baseline expectations for diversifying immune investment, we next wanted to determine what, if any, effect a decoupling cost may have on dTCR investment. We saw that adding a decoupling cost that scaled with the degree of the discrepancy between adult and larval investment significantly altered both larval and adult immune investment (fig. 2E,F). The resulting investment plots resembled an enhanced version of the larval investment predictions (fig. 2C). Adult dTCR investment was quantitatively (but not qualitatively) diminished compared to the case with no decoupling costs (fig. 2D). Larvae no longer invested when adult parasite diversity was less than iTCR repertoire diversity but invested substantially more when adult parasite diversity exceeded iTCR diversity. Adult dTCR investment decreased faster along the dTCR cost axis than in the uncoupled case but remained high when parasite diversity was high and dTCR costs were low. These results suggest that decoupling costs have the potential to disrupt optimal dTCR immune strategies across both life stages.

### Larval dTCR investment depends on the length of the larval life stage and larval parasite diversity

Expanding the results from fig. 2, we next wanted to evaluate whether host dTCR investment was dependent on larval rather than adult parasite diversity. We predicted that larval stage dTCR investment would increase with parasite diversity and decrease with dTCR cost, mirroring adult optima (Fig. 2). We also wanted to determine how varying the length of the larval stage would affect larval dTCR investment, so we conducted simulations with standard (25% of lifespan) and long (75% of lifespan) larval stage lengths. Between these two conditions, we expected that the long larval stage simulations would exhibit an investment pattern that was closer to the adult investment pattern in fig. 2D while the short larval stage would be more likely to exhibit a novel investment pattern. We further predicted that as larval parasite diversity increased, adult investment in dTCR based immunity would be largely invariant because adult stage parasite diversity was high in all simulations.

In simulations with short larval stages, larvae did not invest in dTCR immunity until larval parasite diversity neared its maximum, and was highest when dTCR costs were intermediate. Larval investment in dTCR immunity in these simulations never reached the levels seen in hosts in other conditions (fig. 3A). Simulations with a long larval stage behaved as predicted, with larval dTCR investment increasing as larval parasite diversity exceeded iTCR repertoire diversity (fig. 3C) and decreasing as dTCR costs increased, mimicking the investment pattern observed in adults in fig. 2D.

**Figure 3.**
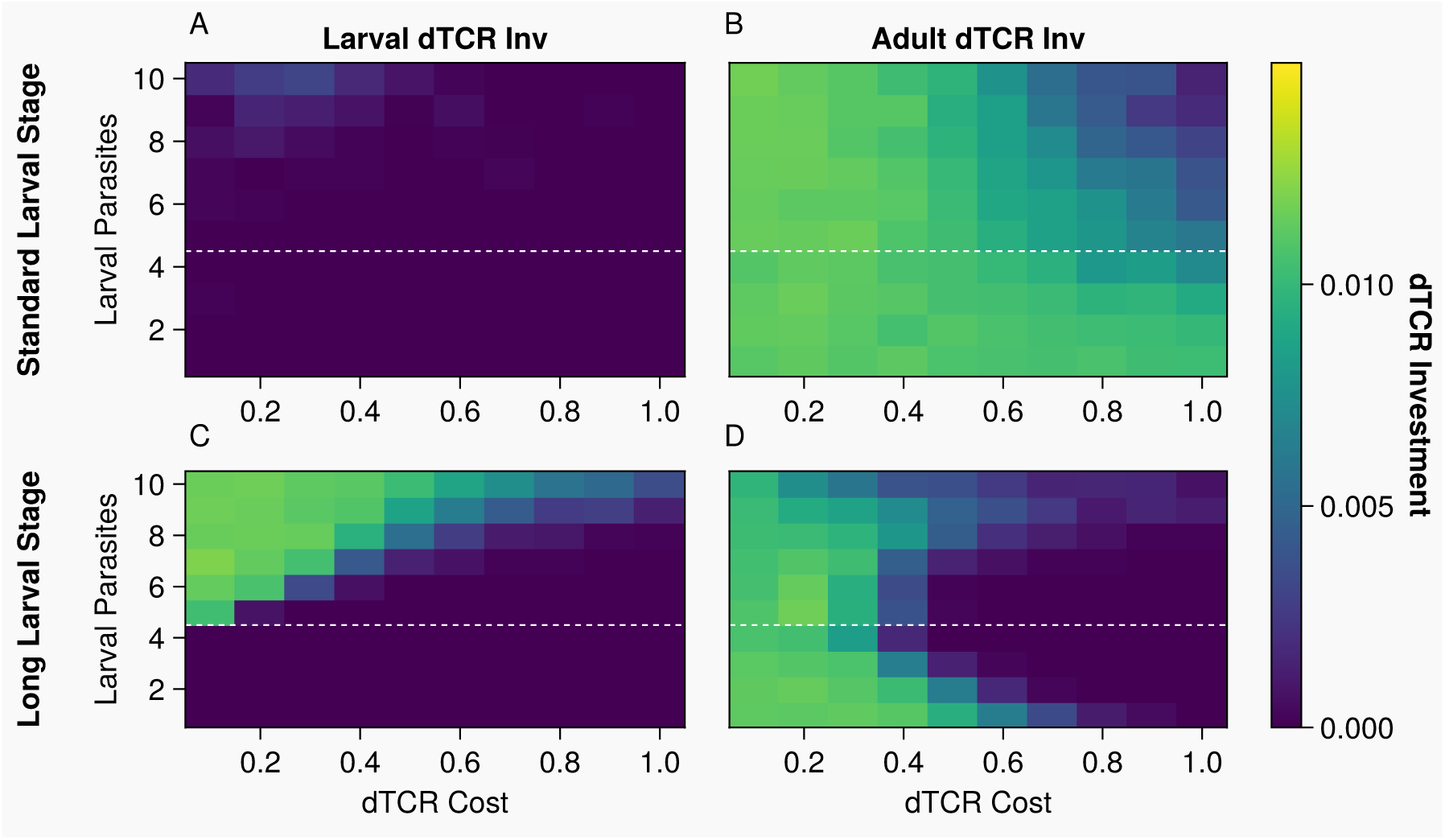
The influence of parasite diversity on dTCR investment depends on stage length. The y-axis shows larval parasite diversity, the x-axis shows dTCR cost. Top row: larval dTCR investment. Bottom row: adult dTCR investment. Left column: hosts spent 75 timesteps as larvae. Right column: hosts spent 25 timesteps as larvae. The dashed white line shows iTCR repertoire diversity.

Adult dTCR investment also changed significantly with larval stage length (fig. 3B,D). In these simulations, adults were always exposed to the maximal parasite diversity. In long larval life stage simulations, adult dTCR investment followed a U-shaped distribution, where dTCR investment was high when the cost was low, and as dTCR costs increased, investment was only high when larval parasite diversity was low or high (fig. 3D). In simulations with short larval life stages, adult dTCR investment more closely resembled the pattern observed in fig. 2D. Moderate decoupling costs altered dTCR investment similarly to the simulations shown in fig. 2, with investment decreasing along the dTCR cost access and increasing along the larval parasite diversity axis (fig. S5).

#### Changes in dTCR investment arise from “all-or-nothing” investment dynamics

In the results so far, dTCR immune investment exhibited a range of intermediate values depending on variation in ontogenic and environmental factors, but it is unclear if these changes arose from a shift in the mean investment or the relative weights of multimodal investment distributions. For example, directional selection might favor differences in mean investment across populations, but diversifying selection might promote coexistence of host genotypes within populations that heavily invest in dTCR immunity with those that do not. We returned to the adult parasite diversity vs. dTCR cost simulations from figure 2D to disentangle these possibilities.

We observed a decrease in mean dTCR investment as dTCR costs increased (fig. 4A). When looking at the mean investment from each of the 100 simulations that contributed to each pixel of the plotted means, very few simulations had a mean dTCR investment between the maximal and minimal values (fig. 4B-D). Rather, most individual simulations converged on two alternate equilibria of “high” (0.01, a value that guarantees the generation of a matching dTCR against any parasite) or “no” investment along the gradient of dTCR costs, and each pixel reflects the investment from individual populations that gravitated toward one extreme or the other.

**Figure 4.**
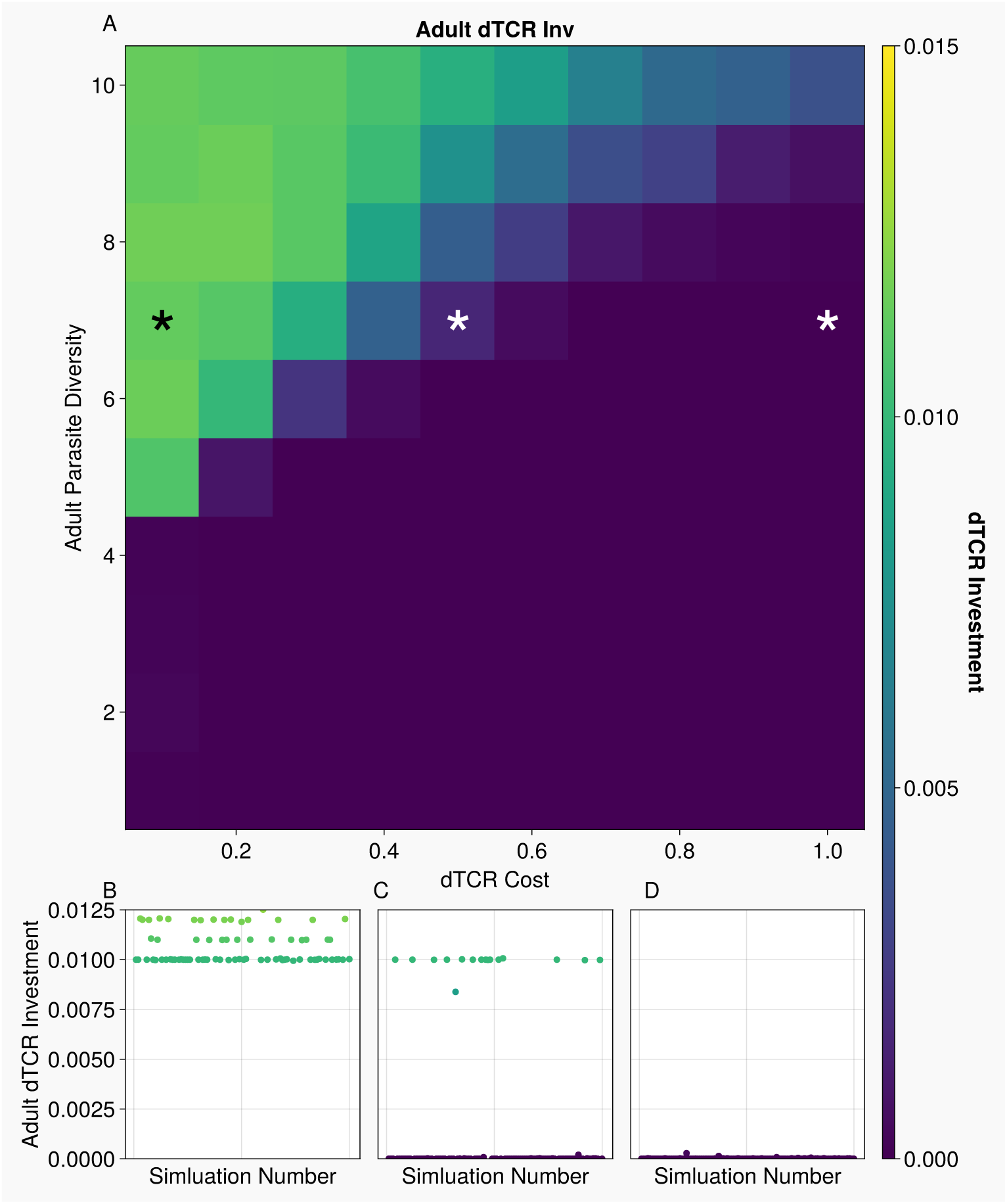
Changes in mean dTCR investment arise from shifts in the proportion of populations that converge on minimal or maximal investment, not a shift in unimodal population mean values. A) adult dTCR investment: the y-axis shows adult parasite diversity, the x-axis shows dTCR cost. Warmer colors indicate greater dTCR investment. The asterisks indicate the conditions that were investigated in detail in panels B,C, and D. B,C,D) simulation mean dTCR investment values when dTCR costs were .1 (B), .5(C), and 1.0 (D). The y-axis shows mean dTCR investment from each simulation that was averaged together for the top plot, separation between points along the x-axis is purely to separate otherwise overlapping data points.

### Intermediate dTCR investment is rare, but not totally absent in our simulations

Populations within our simulations tended to either invest maximally in dTCRs or not at all. However, there were instances where we observed intermediate investment (fig. S6-9). Larval hosts seemed particularly predisposed to intermediate investment in simulations varying dTCR maturation time against the percent of life spent as a larva (figs. 5D, S7). We wanted to determine if these values represented true intermediate investment or just an unusually slow convergence on one of the extreme equilibria.

In simulations where dTCR maturation took longer than the larval life stage (asterisk, fig. 5A,C) larvae did not evolve investment in dTCR immunity in the absence of a decoupling cost (fig. 5B), but some populations of hosts (17/100 simulations) evolved intermediate investment in larval dTCR immunity (between .005 and .01) in simulations with moderate decoupling costs (fig. 5D). This behavior was stable for hundreds of generations (fig. 5D) and was not just a delay in time to fixation.

**Figure 5:**
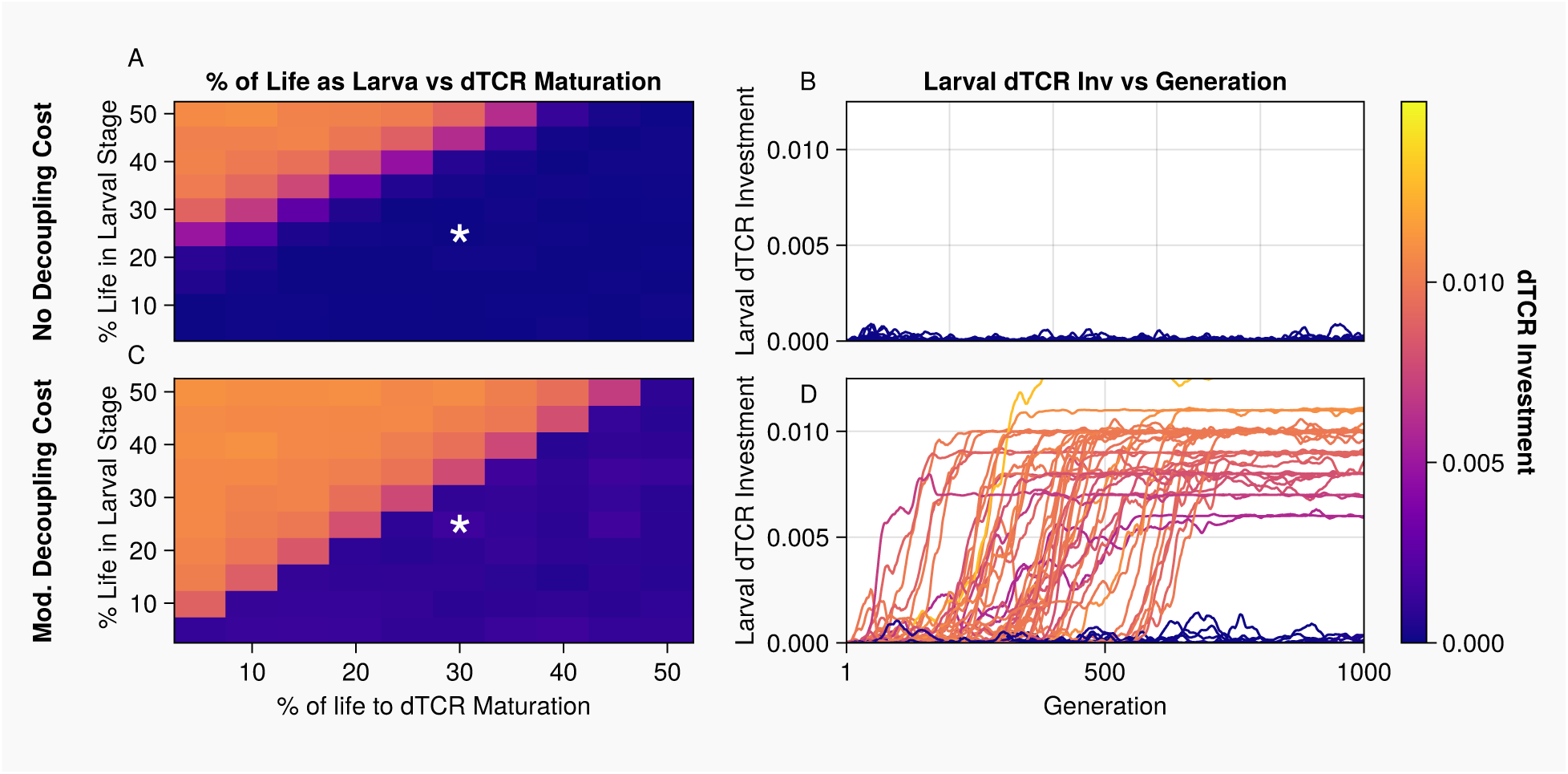
Under certain conditions larvae exhibit intermediate dTCR investment strategies. A,C) larval stage dTCR investment when varying % of life in larval stage (y-axis) vs time taken for dTCR maturation. B,D) Mean larval dTCR investment in each population across generations from the simulations where larval life-stage was 25 timesteps and dTCR maturation took 30 timesteps, y-axis larval dTCR investment and x-axis shows generations. There was no decoupling cost in A and B, and an intermediate decoupling cost in C and D. Despite a number of simulations reaching moderate or high dTCR investment, more than 80% of simulations still had zero or near zero dTCR investment. Warmer colors indicate greater dTCR investment.

## Discussion

We built an evolutionary model of stage-structured dTCR investment over ontogeny to clarify when patterns of investment in TCR diversity reflect classical predictions for innate and adaptive immunity and when we need to consider the specific qualities of T cells in refining these predictions. As with traditional models of adaptive immune system evolution, dTCRs were more likely to evolve on top of iTCRs when the costs associated with them were low and parasite diversity was high (Mayer et al. 2016). dTCR investment was also more likely in life stages that were longer and more likely to experience infection by novel parasites, as predicted by recent theory on specific immune investment (Downie et al. 2021). The temporal lag required for dTCR maturation and selection is a rarely considered cost, however, and here we show that it interacts with stage length to dampen dTCR investment where it might otherwise be favorable (fig. S2). Moreover, we show that the cost of decoupling immune systems across stages, which may be disproportionately genetic or physiological in non-metamorphic species and resource-based in metamorphic species (Urlacher et al. 2018; Collet et al. 2023), substantially dampens investment away from otherwise optimal levels in both stages. Finally, our model suggests that dTCR investment is usually an all-or-nothing gambit. Just as the tiny minifish (*Paedocypris sp*.) invests in dTCRs at rates equivalent to mammals (Giorgetti et al. 2021) even though it can only physically fit about 10,000 T cells in its body, our *in silico* populations evolved either maximal or no investment in dTCRs under most scenarios. The one exception is when organisms incur decoupling costs, in which case larvae sometimes invest at intermediate levels, presumably to minimize the fitness costs associated with decoupled immune investment strategies. The predictions from this model should be directly testable in real-world populations of amphibians, where we expect to see a lower probability of larval dTCR investment for species that have short larval durations relative to adult stages and higher larval parasite burdens (Table 1). For those species that do invest in dTCRs as larvae, they should have repertoires of equivalent diversity to adults.

A key result of our model was the presence of selection on dTCR investment driving entire populations to either invest highly in dTCR immunity or rely solely on iTCR immunity for defense against parasites (fig. 4). At intermediate parasite diversity and dTCR cost levels, this led to an intermediate frequency in larval dTCR investment across populations (fig. 5). This means that if one were to sample several ponds across a frog species’ range, some may sport tadpoles that have a dependable dTCR repertoire while others may sport only iTCRs. Because there is not an abundance of data on the diversity of parasites across stages for most anurans (Bienentreu and Lesbarrères 2020) the microbial diversity data in Table 1 also includes commensal microbiota and is a questionable proxy for this metric. In a similar vein, it is hard to quantify variation in the cost of dTCR investment. Physiological proxies for these costs, such as thymus size and its variation across species, could serve as a stand in for investment capacity (Miodoński et al. 1996; Gui et al. 2012). We conducted a preliminary analysis of the relationship between general microbial diversity and stage length (supplemental methods and results), where we found a weak but significant relationship between relative stage length and microbial diversity (fig. S3).

Our results predict that decoupling costs play a critical role in driving dTCR investment across both the larval and adult life-stages (figs. 2, S1,S2). Moderate and high decoupling costs reshaped both larval and adult dTCR investment, generally expanding the conditions under which larvae invested in dTCR immunity and reducing the conditions in which adults invested. Decoupling costs could thus provide a biological explanation for differences between observed immune strategies and theoretically optimal ones. Some species of amphibians may leverage metamorphosis as a tool to compensate for such decoupling costs, as they only mount competent immune responses to specific pathogens following metamorphosis (Basanta et al. 2023). To the extent that complete metamorphosis could reduce overall decoupling costs by allowing organisms to optimize for either life stage through extensive body plan remodeling (Mcmahon and Hayward 2016), we might expect metamorphic anuran species to show greater variance in dTCR investment across stages than direct-developing species, which may have shallower differences across ontogeny but lower investment overall.

Our results also highlight the potential for cross-stage dTCR immunity, where larval dTCR investment is used to defend adults that do not invest in dTCR immunity, as a viable strategy. For example, we found that larval dTCR investment increased as dTCR cost increased under some conditions, especially just after parasite diversity exceeded iTCR repertoire diversity (figs. 2C,3B). When investigating this unintuitive behavior, we noticed that under these specific conditions, hosts were primarily investing in dTCR immunity in either the adult or larval stage but very rarely in both stages (fig. S10). At lower dTCR costs dTCR investment was most common in adults, but as dTCR costs increased investment shifted to the larval stage until dTCR costs became too high and investment stopped entirely. Our interpretation of this pattern is that the shorter larval stage length curtails the accumulation of costs from investing in dTCR immunity, while at the same time providing a decent probability of generating dTCRs that would be useful later in life. In the wild, larval immune memory has been shown to persist across metamorphosis (Robert et al. 1997), suggesting that coasting into adulthood on larval dTCRs could be a viable real-world immune strategy.

Amphibians are not the only stage-structured organisms for which this model might be relevant. Insects do not possess adaptive immune responses, but they do undergo various forms of complete and incomplete metamorphosis and rely on innate immune signaling pathways that function pleiotropically with development (Williams et al. 2023). Thus, they may represent a good model for investigating decoupling costs associated with immune investment across life stages or to understand variance in immune priming, a form of innate immune memory in insects (Prakash and Khan 2022). By the same token, plants have stage-structured life histories, and a strong combination of data and theory has provided new insight into the evolution of juvenile susceptibility against plant pathogens (Bruns et al. 2022). Critically, this work found that disease resistance was modulated by different mechanisms between adult and juvenile stages, with later work demonstrating that the fitness costs associated with immune defense also change with age (Slowinski et al. 2025). Furthermore, juvenile resistance can reduce adult reproductive fitness, suggesting a trade-off between survival and reproduction (Buckingham et al. 2023). The relationship between host life stage and resistance also directs pathogen evolution (Bruns et al. 2012), suggesting future modeling work could take co-evolutionary forces into account to better understand stage structure and immunity.

In modeling studies there is a necessary trade-off associated with model complexity: simple models are easier to interpret while complex models are better able to capture the real world. This work required several simplifying assumptions that may serve as fertile ground for future modeling efforts. For example, our model employed synchronous generations. While we did not include epidemiological dynamics in this model, asymmetry in susceptibility and transmission among stages and individuals would likely have important effects on simulation outcomes (Portner et al. 2022) and increase dependence on the synchrony assumption. Another consideration is the relationship between parasite burden and larval mortality. While there is evidence for parasites having an extreme fitness burden on frogs peri- and post-metamorphosis (Goodman and Johnson 2011), infected larvae are not completely incapable of surviving to sexual maturity and breeding. Our stringent gating of tadpole infection leading to mortality may thus favor an unrealistic degree of larval stage immune competence. Finally, we used fixed cost structures to evaluate host fitness and did not let those evolve to become less costly, as might be the case for a mechanism that increases peripheral tolerance to dTCR pathology, for example. Future modeling work could expand on our findings by building more detailed models of specific life history traits and immune processes of interest, treating this model as a general framework.

Ontogenic and environmental factors like resource availability (Lee et al. 2008; Boots 2011), life-stage (McNeil et al. 2010), and individual immune biases (Howard et al. 2022; Kirouac et al. 2023) shape the immune responses of not just vertebrates, but all organisms with immune systems. Our results provide predictions for expected dTCR investment strategies in wild amphibian populations as well as the environmental and life history conditions that could give rise to rare but important exceptions to the rule, like those that moderately invest in dTCR immunity. Given the worldwide amphibian biodiversity crisis linked to pathogen susceptibility (Scheele et al. 2019), understanding how life history and ecology interact to define immune investment strategies in amphibians is an urgent need. More generally, these results invite new hypotheses on the evolution of metamorphosis and the nature of decoupling costs associated with stage-structured life histories.

## Supporting information

Supplemental materials for the associated manuscript

## Acknowledgments

We would like to thank Veronica Urgiles for conducting and interpreting the phylogenetic generalized least squares analysis in the supplemental results.

## Data & Code Availability

Data, Code, and a comprehensive ReadMe are available at the following DOI: [when we are happy with figures I will upload to Dryad or Zenodo, depending on cost]

## Supplemental Materials

**Supplemental Figure 1:** dTCR investment generally increases with higher parasite diversity and lower costs when hosts begin with intermediate dTCR investment.

**Supplemental Figure 2:** dTCR investment generally increases with higher parasite diversity and lower costs when hosts begin with intermediate dTCR high.

**Supplemental Figure 3:** Loess plot showing the phylogenetically-corrected relationship between tadpole stage length and the ratio of larval to adult stage microbial diversity.

**Supplemental Figure 4:** The delay between infection and dTCR activation acts like extended dTCR maturation time.

**Supplemental Figure 5:** The influence of parasite diversity on dTCR investment depends on stage length.

**Supplemental Figure 6:** In simulations varying the adult parasite diversity against dTCR costs there are few intermediate investors.

**Supplemental Figure 7:** In simulations varying the percent of life spent as a larva against dTCR costs a large number of simulations have intermediate investing larvae.

**Supplemental Figure 8:** In simulations varying the larval parasite diversity against dTCR costs, when the larval life stage was 25 timesteps, there are few intermediate investors.

**Supplemental Figure 9:** In simulations varying the larval parasite diversity against dTCR costs, when the larval life stage was 75 timesteps, there are few intermediate investors.

**Supplemental Figure 10:** Larval dTCR investment vs Adult dTCR investment when adult parasite diversity was 5, dTCR cost was .2, and there was no decoupling cost.

