## Supplemental materials for the associated manuscript for "The evolution of investment in innate-like and diversified T cell receptors across development"

### Supplemental Methods and Results

We used the studies presented in Table 1 (main text) to analyze the relationship between amphibian larval stage length and microbial diversity in the larval relative to adult stage. We recognize that total microbial diversity may not reflect the quantity of pathogenic microbes the immune system is challenged with. However, such data are lacking in the literature, especially for larval stages. For example, a review of over 800 amphibian pathogen studies conducted between 2009-2019 found that 78% only investigated chytrid fungi or ranaviruses, and only 28% included any larval-stage hosts (Bienentreu and Lesbarrères 2020). Thus, we used total microbial diversity as a potential rough approximation of higher or lower levels of pathogenic microbes, and we acknowledge that validation of this relationship is needed. Data on tadpole and adult stage durations were compiled from AmphibiaWeb (https://amphibiaweb.org/).

Each of the studies included in Table 1 measured skin or gut microbial diversity from both larval and adult samples collected from the same species, location, and time point. However, because these studies used a range of metrics and methodologies to estimate microbial diversity, we could only compare them using the ratio of larval to adult microbial diversity, and could not directly compare larval duration with larval microbial diversity.

We used the *caper* package (Orme et al. 2023) to conduct Phylogenetic Generalized Least Squares (PGLS) regressions in R. We log-transformed the tadpole duration data to normalize it, and used the Pyron and Wiens (2011) pruned amphibian phylogeny to account for shared evolutionary history across species. Because visual inspection of the data suggested a non-linear relationship, we ran a linear model (pgls:Microbial_ratio ~ log_tadpole_duration) as well as a model with a quadratic term (pgls;Microbial_ratio ~ log_tadpole_duration_lin + log_tadpole_duration_quad; Zagmasjster et al. 2014). The linear model showed a non-significant association ($p = 0.21$) with low $R^2$ (~0.11). The model including a quadratic term (Supplemental Figure 10) showed a slightly significant linear term (estimate = –2.52, $p = 0.045$), a slightly non-significant quadratic term (estimate = –2.68, $p = 0.062$), and a modest relationship ($R^2 = 0.28$). However, the linear-only model AIC score was 2.4 lower than the model including the quadratic term, suggesting the linear model is a better fit.

The small number of studies we found, lack of direct pathogenic diversity estimation, and weak relationships preclude drawing robust conclusions from this analysis. However, we find the patterns suggestive of potential relationships, and encourage more research into pathogenic communities in larval amphibians.

**Table of parameters**

| **Biological Feature** | **Model parameter name** | **Default** | **Biological Explanation** |
|---|---|---|---|
| Population Size | numHosts | 500 | Simulation population size |
| Number of Generations | Generations | 1000 | Number of generations simulated |
| Adolescent life-stage length | perAdol | .25 | Percent of life spent as an adolescent (note, .25*100 = 25 timestep stage length) |
| Host lifespan | lifespan | 100 | Total number of timesteps between birth and death/reproduction |
| dTCR generation cost | genCost | .01 | dTCR investment level necessary to generate a perfect fit dTCR |
| Innate-like TCR repertoire size | numiTCR | 4 | Initial iTCR repertoire size, |
| Initial iTCR match against parasites | perMatch | 100 | Percent of a parasite that initial iTCRs are guaranteed to match |
| Larval parasite diversity | numTadPars | 7 | Number of larval parasite species |
| Adult parasite diversity | numAdulpars | 10 | Number of adult parasite species |
| Parasite complexity | parLen | 20 | Number of bases each parasite is composed of, longer parasites are more complex |
| Larval infection chance | adolInfMod | 1.0 | Modifier for larval infection chance based on adult infection chance, 1.0 means identical infection chance |
| Host infection chance | infChan | .05 | Chance of an infection event occurring in a given timestep |
| Cost of immune action | immCost | 1 | Modifier for the cost of mounting an immune defense |
| Cost of infection/parasite burden | parCost | 1 | Modifier for the cost of parasite burden |
| Cost of iTCR use | iTCRCost | .1 | Fitness cost associated with using an iTCR |
| Cost of dTCR use | dTCRCost | .1 | Fitness cost associated with using a dTCR |
| Decoupling Cost | stageCost | 0,1,10 | Fitness cost for differential adult and larval dTCR investment |
| dTCR maturation time | dTCRMat | 15 | Timesteps before dTCR immune response matured. |

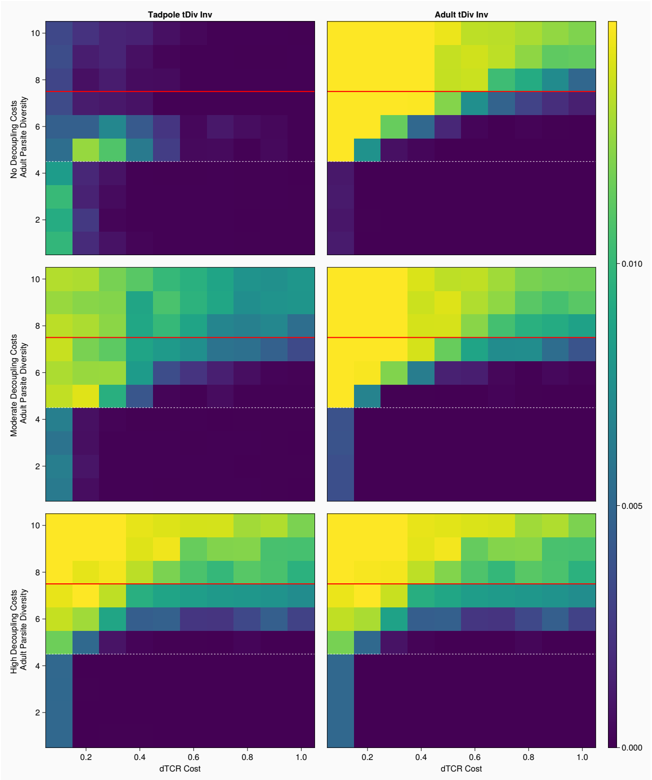


**Supplemental Figure 1:** dTCR investment generally increases with higher parasite diversity and lower costs when hosts begin with intermediate dTCR investment. Y-axis shows adult parasite diversity, x-axis shows the cost associated with generating dTCRs. Top row: no decoupling cost, middle row: moderate decoupling cost, bottom row: high decoupling cost. Warmer colors indicate increased investment.

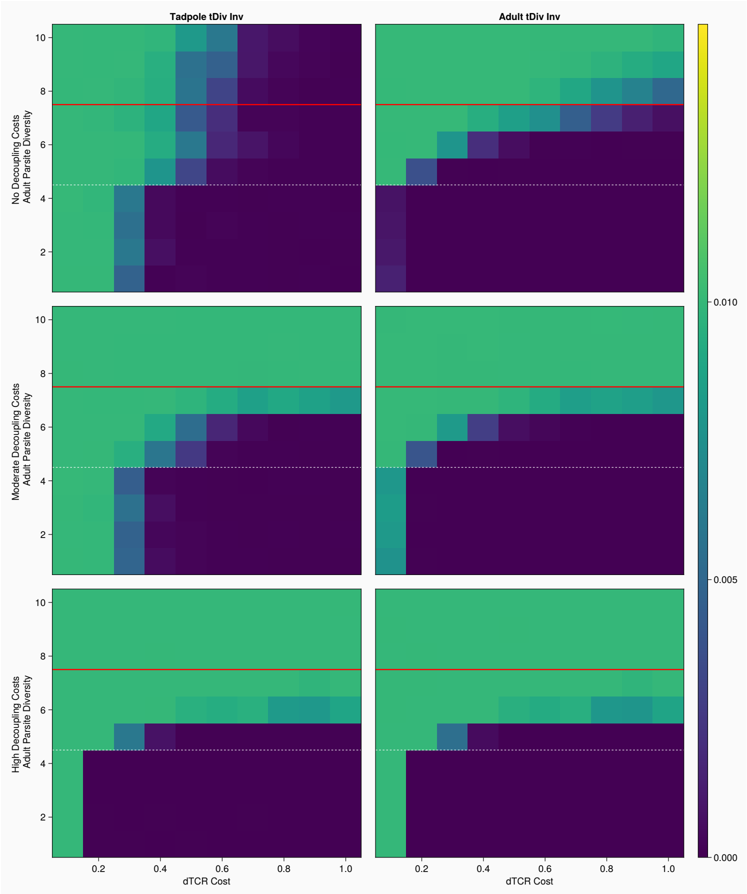


**Supplemental Figure 2:** dTCR investment generally increases with higher parasite diversity and lower costs when hosts begin with intermediate dTCR high. Y-axis shows adult parasite diversity, x-axis shows the cost associated with generating dTCRs. Top row: no decoupling cost, middle row: moderate decoupling cost, bottom row: high decoupling cost. Warmer colors indicate increased investment.

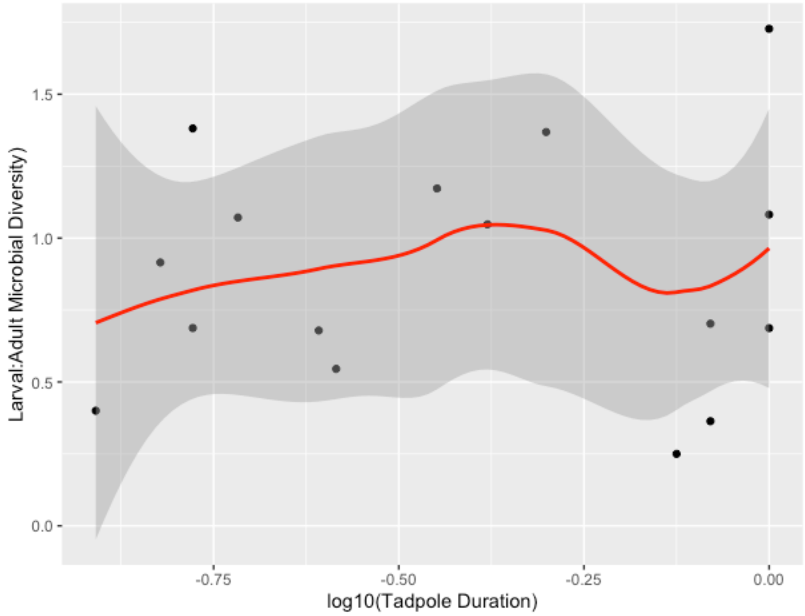


**Supplemental Figure 3:** Loess plot showing the phylogenetically corrected relationship between tadpole stage length and the ratio of larval to adult stage microbial diversity. Data were estimated from published studies that quantified larval and adult microbiome diversity for the same species, location, and sampling date.

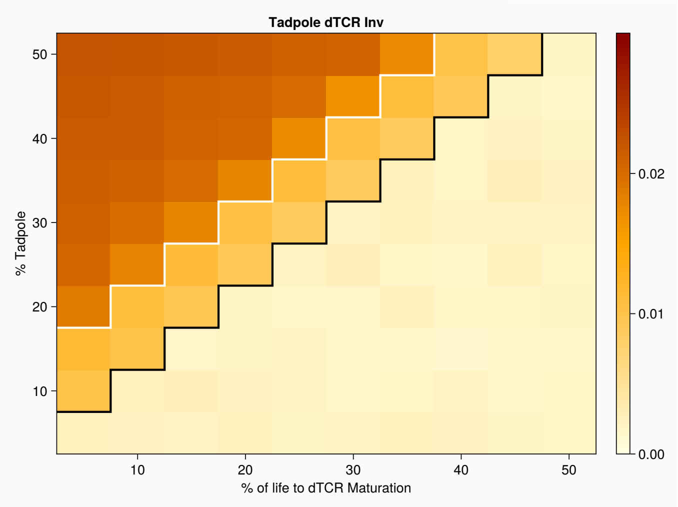


**Supplemental Figure 4:** The delay between infection and dTCR activation acts like extended dTCR maturation time. y-axis shows the percent of life in the larval stage, x-axis shows the percent of life before the diversifying immune response matures. The white line separates the area where both sets of larvae were investing in immunity from the area where just larvae that evolved without a dTCR delay were investing in immunity. The black line separates the area where the larvae that evolved without a dTCR delay were investing in immunity from the area where no larvae invested in the dTCR immune response. Darker colors indicate greater dTCR investment.

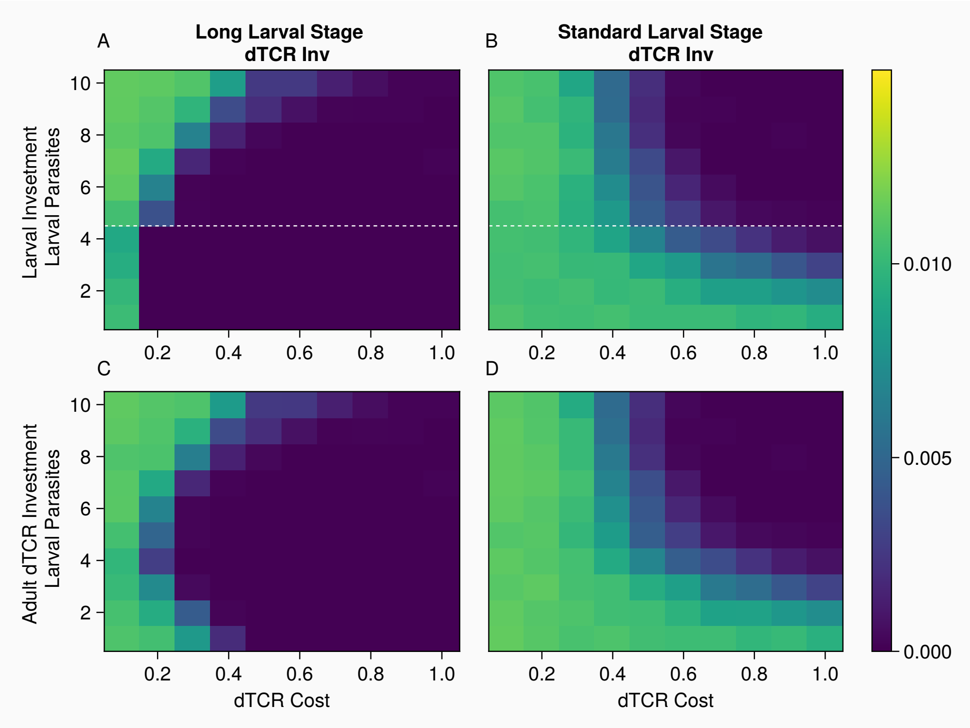

**Supplemental Figure 5:** The influence of parasite diversity on dTCR investment depends on stage length. The y-axis shows larval parasite diversity, the x-axis shows dTCR cost. The left plot shows simulations where hosts spent 75 timesteps as larvae, the right plot shows simulations where hosts spent 25 timesteps as larvae. The dashed white line shows iTCR repertoire diversity.

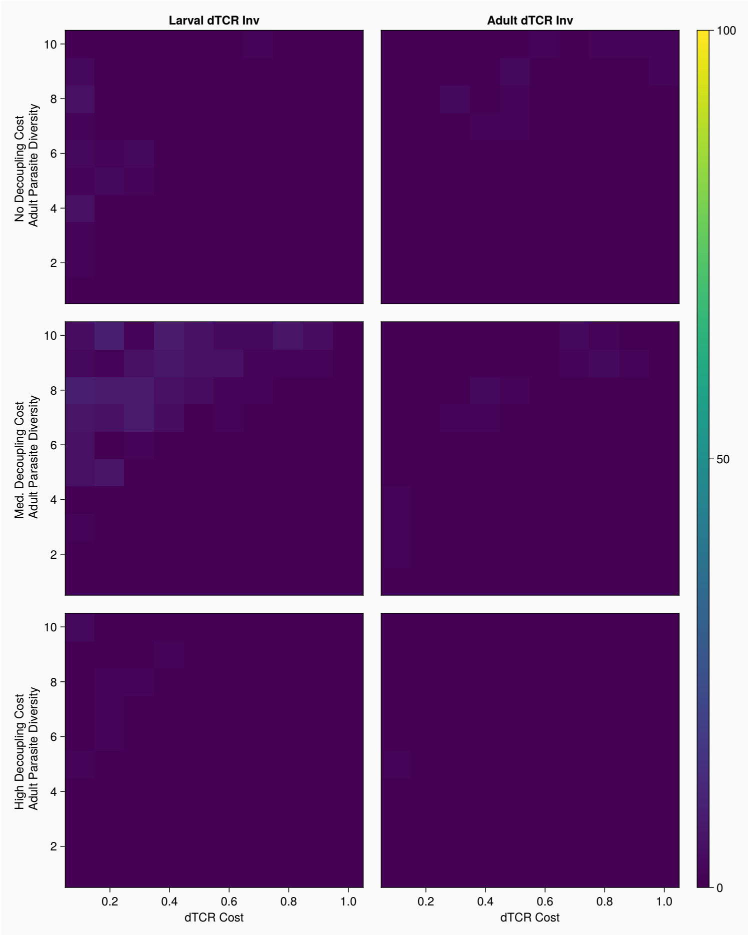


**Supplemental Figure 6:** In simulations varying the adult parasite diversity against dTCR costs there are few intermediate investors. The y-axis shows adult parasite diversity, the x-axis shows dTCR cost. Top row: no decoupling cost, middle row: moderate decoupling cost, bottom row: high decoupling cost. Warmer colors indicate a greater number of simulations with intermediate-investing hosts.

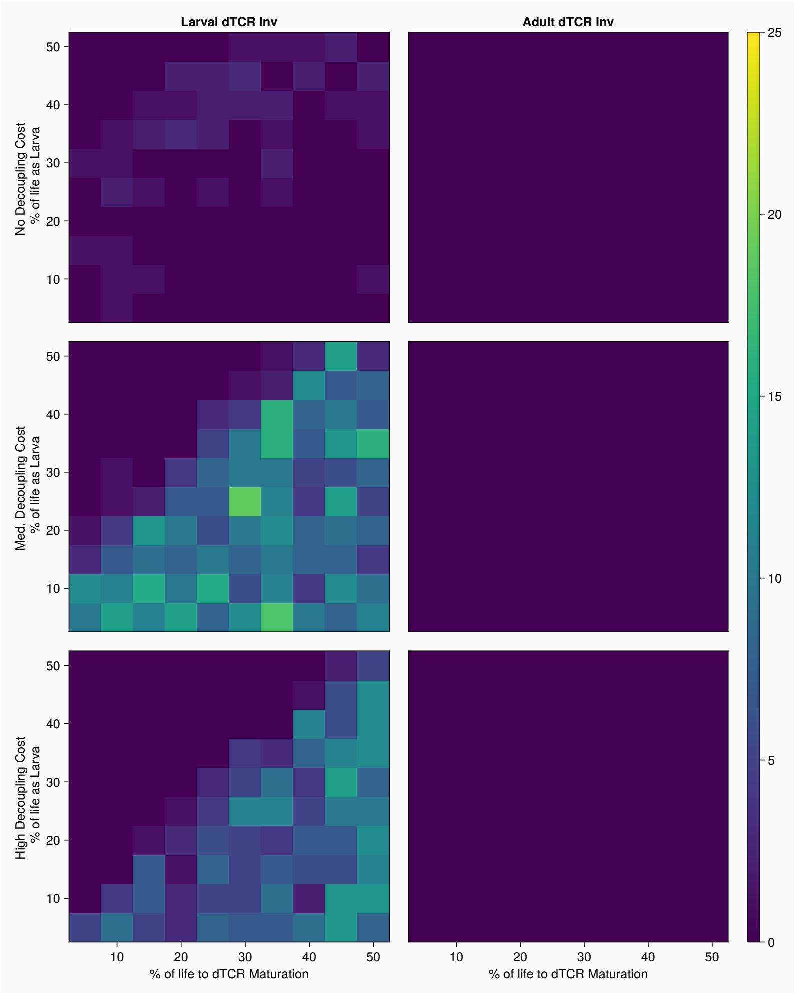


**Supplemental Figure 7:** In simulations varying the percent of life spent as a larva against dTCR costs a large number of simulations have intermediate investing larvae. The y-axis shows the percent of life spent as a larva; the x-axis shows dTCR cost. Top row: no decoupling cost, middle row: moderate decoupling cost, bottom row: high decoupling cost. Warmer colors indicate a greater number of simulations with intermediate investing hosts.

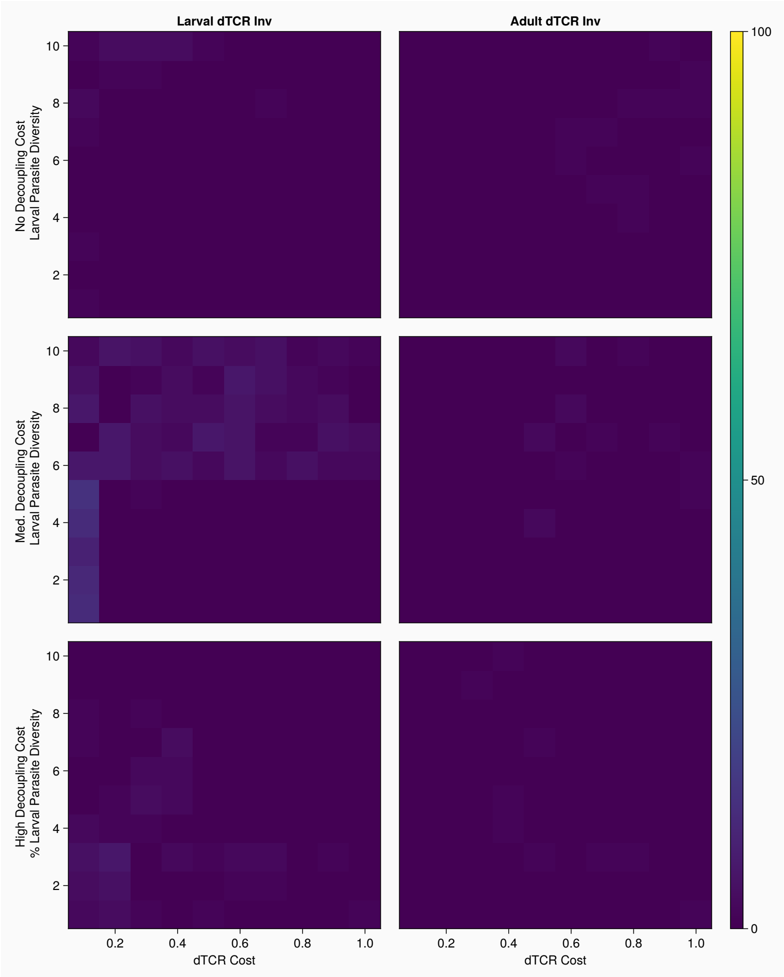


**Supplemental Figure 8:** In simulations varying the larval parasite diversity against dTCR costs, when the larval life stage was 25 timesteps, there are few intermediate investors. The y-axis shows larval parasite diversity, the x-axis shows dTCR cost. Top row: no decoupling cost, middle row: moderate decoupling cost, bottom row: high decoupling cost. Warmer colors indicate a greater number of simulations with intermediate investing hosts.

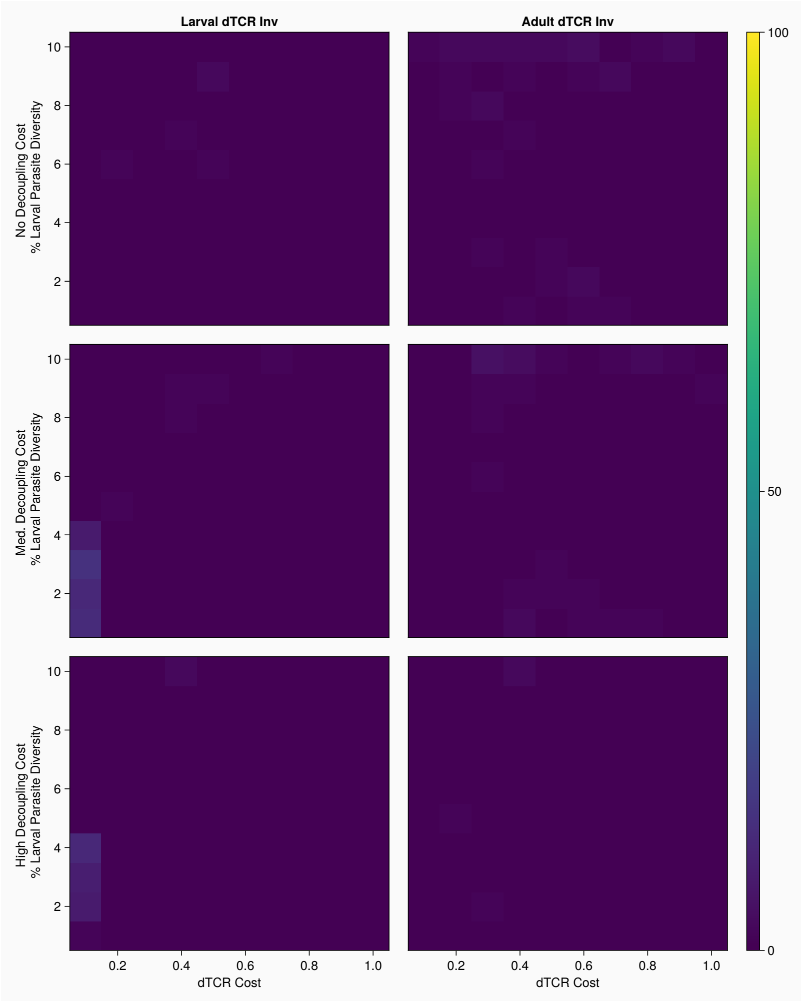


**Supplemental Figure 8:** In simulations varying the larval parasite diversity against dTCR costs, when the larval life stage was 75 timesteps, there are few intermediate investors. The y-axis shows larval parasite diversity, the x-axis shows dTCR cost. Top row: no decoupling cost, middle row: moderate decoupling cost, bottom row: high decoupling cost. Warmer colors indicate a greater number of simulations with intermediate investing hosts.

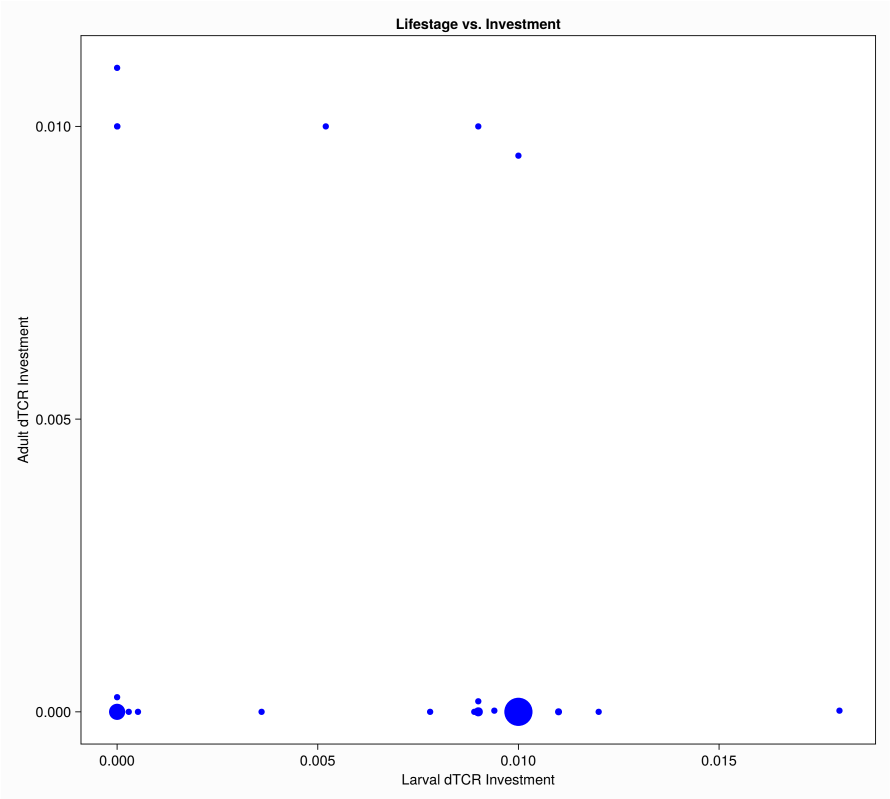


**Supplemental Figure 10:** Larval dTCR investment vs Adult dTCR investment when adult parasite diversity was 5, dTCR cost was .2, and there was no decoupling cost. Each dot comes from a single generation, X-axis shows the larval dTCR investment level; y-axis shows adult dTCR investment. The size of each dot shows the number of simulations with that combination of (Larval investment, adult investment).
